# CRISPR-mediated engineering of bovine satellite cells for Alpha-Gal Syndrome-compatible cultivated meat

**DOI:** 10.64898/2026.05.20.726299

**Authors:** Susan D’Costa, Shailesh K. Choudhary, Grace E. Kenney, Jacqueline Shine, Brian Diekman, Scott P. Commins, Douglas H. Phanstiel

**Affiliations:** Thurston Arthritis Research Center, University of North Carolina, Chapel Hill, NC, United States of America; Division of Rheumatology, Allergy and Immunology, University of North Carolina, Chapel Hill, NC, United States of America; Curriculum in Bioinformatics and Computational Biology, University of North Carolina, Chapel Hill, NC, United States of America; Lampe Joint Department of Biomedical Engineering, University of North Carolina, Chapel Hill, NC, United States of America; Institute for Global Health and Infectious Diseases, Chapel Hill, NC, United States of America; Lineberger Comprehensive Cancer Center, University of North Carolina, Chapel Hill, NC, United States of America; Curriculum in Genetics and Molecular Biology, University of North Carolina, Chapel Hill, NC, United States of America; Department of Cell Biology and Physiology, University of North Carolina, Chapel Hill, NC, United States of America

## Abstract

Alpha-gal Syndrome (AGS) is a potentially life-threatening allergy caused by an IgE-mediated immune response to galactose-α-1,3-galactose (alpha-gal), a carbohydrate epitope present in most mammalian meats. Currently, strict avoidance of mammalian meat remains the primary management strategy for affected individuals, and alpha-gal-free beef is not commercially available. Here, we leverage cultivated meat as a biotechnology platform to address this unmet clinical need by engineering alpha-gal–free bovine muscle cells. Using CRISPR/Cas9 genome editing, we disrupted GGTA1, the gene encoding α1,3-galactosyltransferase, in immortalized bovine satellite cells (iBSCs). High-efficiency editing produced clonal GGTA1 knockout iBSCs harboring a homozygous frameshift mutation. Flow cytometry and immunofluorescence confirmed loss of the alpha-gal epitope, while bulk RNA-seq indicated minimal disruption of global gene expression and preserved myogenic differentiation capacity. Importantly, lysates from GGTA1 knockout iBSCs elicited substantially reduced basophil activation in assays using plasma from a patient with AGS, indicating reduced basophil activation consistent with reduced allergenic potential. Together, these findings establish a proof of concept for engineering AGS-compatible cultivated meat and demonstrate the potential of cultivated meat technologies to address human health challenges.

**HIGHLIGHTS:**

∘ CRISPR/Cas9-mediated disruption of GGTA1 eliminated alpha-gal from bovine satellite cells
∘ GGTA1 knockout cells retained myogenic identity and differentiation capacity
∘ GGTA1 knockout reduced basophil activation in an alpha-gal syndrome immune assay
∘ Genome-edited bovine cells provide a proof of concept for AGS-compatible cultivated meat

## Introduction

**A**lpha-gal Syndrome (AGS) is a severe allergic condition caused by an immunoglobulin E (IgE)-mediated reaction to galactose-α-1,3-galactose (alpha-gal), a carbohydrate epitope present on glycoproteins and glycolipids in most mammalian meat and derived products. Humans lack the ability to synthesize alpha-gal due to evolutionary inactivation of α1,3-galactosyltransferase (α1,3GT), the enzyme responsible for generating the alpha-gal epitope (Galα1,3Galβ1,4GlcNAc-R) (Galili, 2005; Koike et al., 2007). As a result, exposure to alpha-gal can elicit a delayed allergic response in sensitized individuals.

Sensitization typically occurs following tick bites, and subsequent consumption of beef, lamb, or pork can trigger symptoms ranging from gastrointestinal distress and urticaria to life-threatening anaphylaxis (Commins & Platts-Mills, 2013a, 2013b). A fatal case of delayed anaphylaxis following mammalian meat consumption was recently reported, underscoring the potential severity of this condition (Platts-Mills et al., 2025).

Since its initial description in 2009 (Chung et al., 2008; Commins et al., 2009), AGS has been documented on nearly every continent and continues to increase in prevalence, particularly in the southeastern United States where tick exposure is common. From 2010 to 2018, diagnostic testing increased ten-fold while positive test results rose six-fold, and the CDC estimates that up to 450,000 individuals in the United States may have been affected since 2010 (Binder et al., 2021; Thompson et al., 2023). In regions with high tick exposure, alpha-gal has been identified as the leading trigger of anaphylaxis among cases with a definitive cause (Pattanaik et al., 2018), highlighting the growing relevance of this syndrome. Strict avoidance of mammalian meat and derived products remains the primary management strategy for alpha-gal syndrome.

While alpha-gal knockout livestock have been generated experimentally (McGregor et al., 2023; Phelps et al., 2003; Sendai et al., 2006; Tearle et al., 1996; Wei et al., 2022), beef from alpha-gal–deficient animals is not currently commercially available. Cultivated meat offers an alternative approach, enabling the production of muscle tissue from defined cell populations under controlled conditions (Ben-Arye & Levenberg, 2019; Post, 2012; Stephens et al., 2018). This framework also makes it possible to engineer cell lines directly, without requiring generation of whole animals, providing a powerful platform to improve food safety, nutritional properties, and other health-relevant traits. At the same time, recent work comparing cultured primary bovine myoblasts with native tissue found that culture conditions can alter allergen profiles and may increase IgE reactivity to alpha-gal epitopes in cultured cells, highlighting alpha-gal as a specific safety consideration for cultivated beef (Trlin et al., 2026). Targeted disruption of GGTA1, which encodes α1,3-ga-lactosyltransferase, offers a direct strategy to eliminate alpha-gal synthesis and generate bovine muscle cells with reduced AGS-associated immune reactivity.

## Results

### CRISPR-mediated disruption of GGTA1 eliminates alpha-gal in iBSCs

To eliminate alpha-gal production in a cell type relevant to cultivated meat, we used CRISPR/Cas9 to target GGTA1 in immortalized bovine satellite cells (iBSCs), a recently developed cell line with robust proliferative capacity and myogenic potential (Stout et al., 2023). GGTA1 encodes α1,3-galactosyltransferase, the enzyme responsible for alpha-gal synthesis in bovine cells. iBSCs were electroporated with ribonucleoprotein (RNP) complexes containing Cas9 and a guide RNA targeting the fourth exon of GGTA1. Sanger sequencing followed by ICE analysis revealed high editing efficiency in bulk-edited populations, with indel frequencies ranging from 88% to 94% (**Fig. 1A**). Single– base pair insertions were the predominant editing outcome occurring at frequencies of 69% to 85% (**Fig. 1A**). Following clonal isolation and genotyping, three independent clones homozygous for single–base pair insertions were selected for further analysis.

**Fig. 1:**
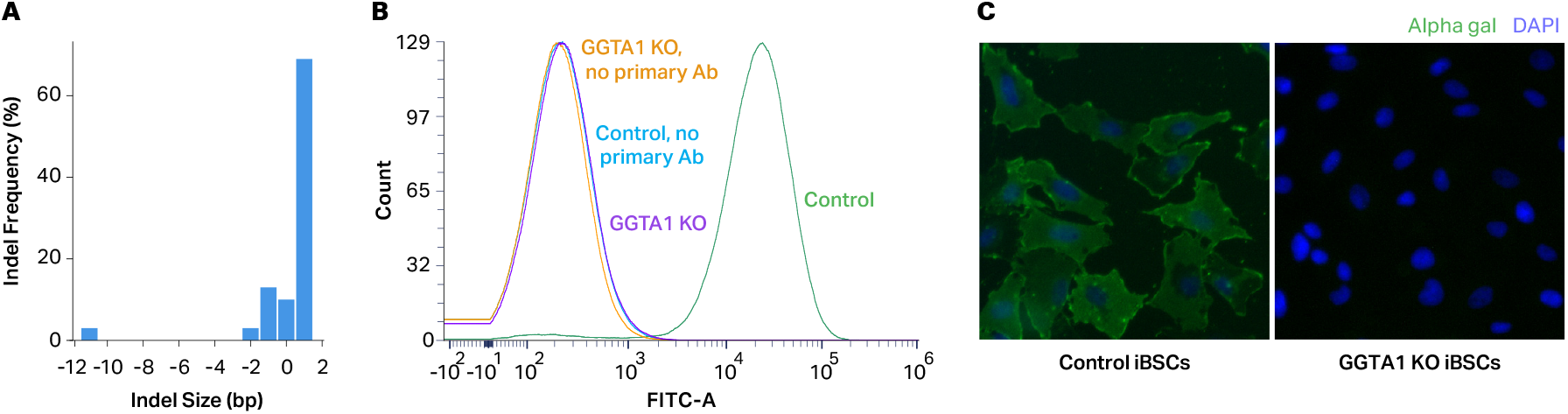
CRISPR-mediated disruption of GGTA1 eliminates alpha-gal in iBSCs. **A**, Predicted relative frequencies of insertions and deletions in bulk-edited iBSCs, as determined by ICE analysis of Sanger sequencing data. **B**, Flow cytometry histograms showing alpha-gal staining in control and GGTA1 KO iBSCs. Cells were stained with an anti–alpha-gal monoclonal antibody followed by a FITC-conjugated secondary antibody. No-primary controls were stained with secondary antibody only. **C**, Immunofluorescence staining showing loss of detectable alpha-gal signal in GGTA1 KO iBSCs. Control and GGTA1 KO cells were stained with an anti–alpha-gal monoclonal antibody and FITC-conjugated secondary antibody, with nuclei counterstained with DAPI.

Alpha-gal expression in edited and untransfected control iBSCs was assessed by flow cytometry and immunofluorescence staining using the alpha-gal– specific monoclonal antibody M86 (**Fig. 1B–C**). Flow cytometric analysis of pooled untransfected control clones (control) showed a clear rightward shift in fluorescence intensity, whereas GGTA1 knockout clones (GGTA1 KO) overlapped with negative control samples, consistent with a complete loss of cell-surface alpha-gal (**Fig. 1B**). Immunofluorescence staining revealed alpha-gal positive cells in control clones but no detectable staining in knockout clones, consistent with elimination of the alpha-gal epitope following GGTA1 disruption (**Fig. 1C**).

### GGTA1 knockout preserves iBSC morphology, differentiation capacity, and overall gene expression profiles

To determine whether GGTA1 disruption altered fundamental properties of iBSCs, we first assessed cellular morphology before and after myogenic differentiation. Phase contrast microscopy revealed that GGTA1 knockout clones were morphologically indistinguishable from unedited control cells, displaying similar cell shape, adherence, and growth characteristics (**Fig. 2A**). Upon induction of differentiation, both control and GGTA1 KO cells exhibited comparable morphological changes consistent with myogenic differentiation, including cell elongation and acquisition of a spindle-shaped phenotype, with no obvious differences between genotypes (**Fig. 2A**).

**Fig. 2:**
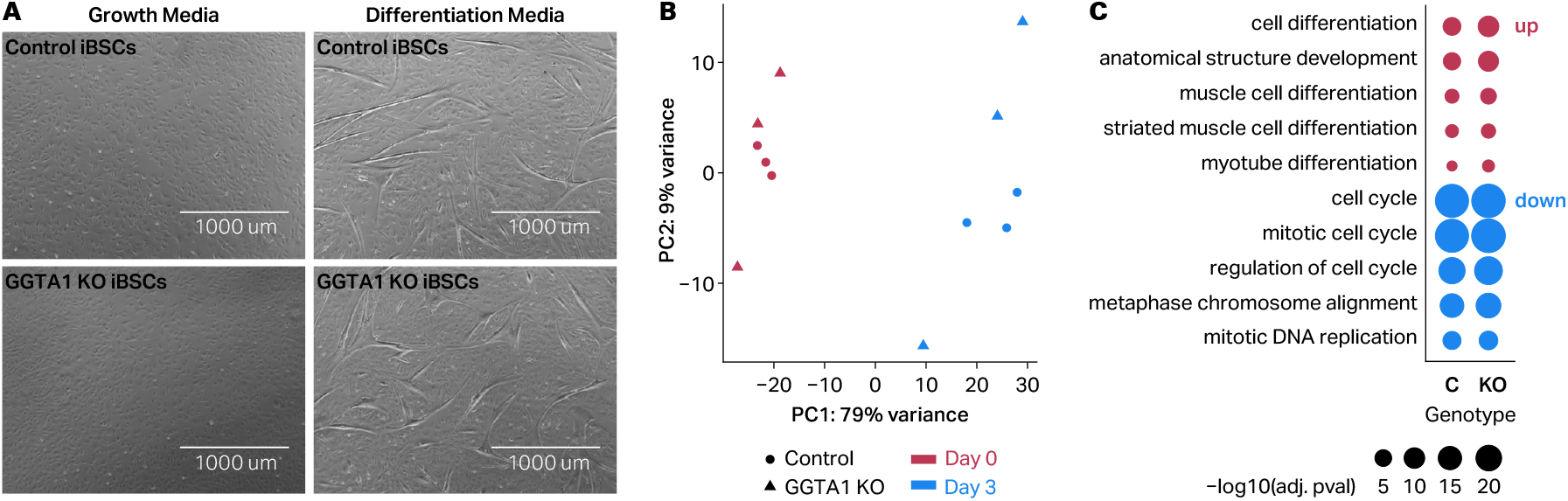
Similar morphological and transcriptomic changes in control and knockout iBSCs during early myogenic differentiation. **A**, Phase contrast images of control and GGTA1 KO iBSCs cultured in growth medium or differentiation medium for five days. **B**, Principal component analysis of RNA-seq data from control and GGTA1 KO iBSCs before differentiation (d0) and after three days in reduced-serum differentiation medium (d3). **C**, Dot plot summarizing selected Gene Ontology terms enriched among genes differentially expressed during differentiation in control and GGTA1 KO iBSCs.

To evaluate the transcriptional consequences of GGTA1 knockout, we performed bulk RNA sequencing on three independent control clones and three independent knockout clones before and after induction of differentiation. Principal component analysis (PCA) demonstrated that the primary source of variation corresponded to differentiation state, with nondifferentiated (d0) samples clustering separately from differentiated (d3) samples along the first principal component (**Fig. 2B**). In contrast, control and GGTA1 KO samples did not segregate by genotype within either state, indicating that GGTA1 disruption did not induce broad shifts in global gene expression.

Differential expression analysis further supported this observation. Induction of differentiation resulted in extensive transcriptional remodeling, with 2,510 and 2,171 genes exhibiting significant changes in control and GGTA1 KO cells respectively (Table S1, DESeq (Love et al., 2014); adjusted p < 0.01, absolute log_2_ fold change > 1). Upregulated genes included regulators of myogenic differentiation and muscle maturation, including IGF2, NFATC4, and CFL2, as well as genes associated with sarcomere assembly and contractile apparatus formation (MYH3, MYBPC2, TNNT2 and TNNC2) and sarcoplasmic calcium homeostasis and muscle contraction (CASQ2 and MYLK). Upregulated genes were enriched for muscle development-related Gene Ontology (GO) terms including muscle cell differentiation (GO:0042692), striated muscle cell differentiation (GO:0051146), and myotube differentiation (GO:0014902) (**Fig. 2C**, Table S2). Down regulated genes were enriched for terms related to cell proliferation (**Fig. 2C**, Table S2). By comparison, only two and thirteen genes met significance thresholds for differential expression between control and GGTA1 KO cells before and after differentiation, respectively (DESeq (Love et al., 2014); adjusted p < 0.01, absolute log_2_ fold change > 1; Table S1). Collectively, these data demonstrate robust activation of the myogenic differentiation program in iBSCs characterized by upregulation of structural, regulatory and contractile genes essential for skeletal muscle formation. Together, these results indicate that GGTA1 KO iBSCs retain morphology, differentiation capacity, and global transcriptional programs comparable to untransfected control cells.

### GGTA1 KO iBSCs induce reduced basophil reactivity

To assess whether elimination of alpha-gal reduced cellular allergenicity, we performed an indirect basophil activation test using lysates from control and GGTA1 KO iBSC clones (Santos & Lack, 2016). Basophils from a healthy donor were sensitized with plasma from an alpha-gal–positive individual and stimulated with cellular lysates. Flow cytometry was used to evaluate basophil activation, activation was quantified as the percentage of CD123^+^ (a basophil marker) cells that were also CD63^+^ (a marker of basophil activation). Negative controls showed minimal activation (≤ 0.38%), whereas positive controls induced robust activation (∼14-21%; **Fig. 3A**), consistent across anti-IgE, cetuximab, and beef thyroglobulin. Lysates from control iBSCs induced basophil activation comparable to alpha-gal–containing positive controls, with a mean activation of 17.93% ± 1.03% (Fig. 3A-B). By comparison, lysates from GGTA1 KO clones produced a marked reduction in basophil activation, averaging 6.40% ± 2.56% (Fig. 3A-B). Although low levels of activation were observed for some clones, responses to knockout lysates were consistently and substantially lower than those elicited by control lysates.

**Fig. 3:**
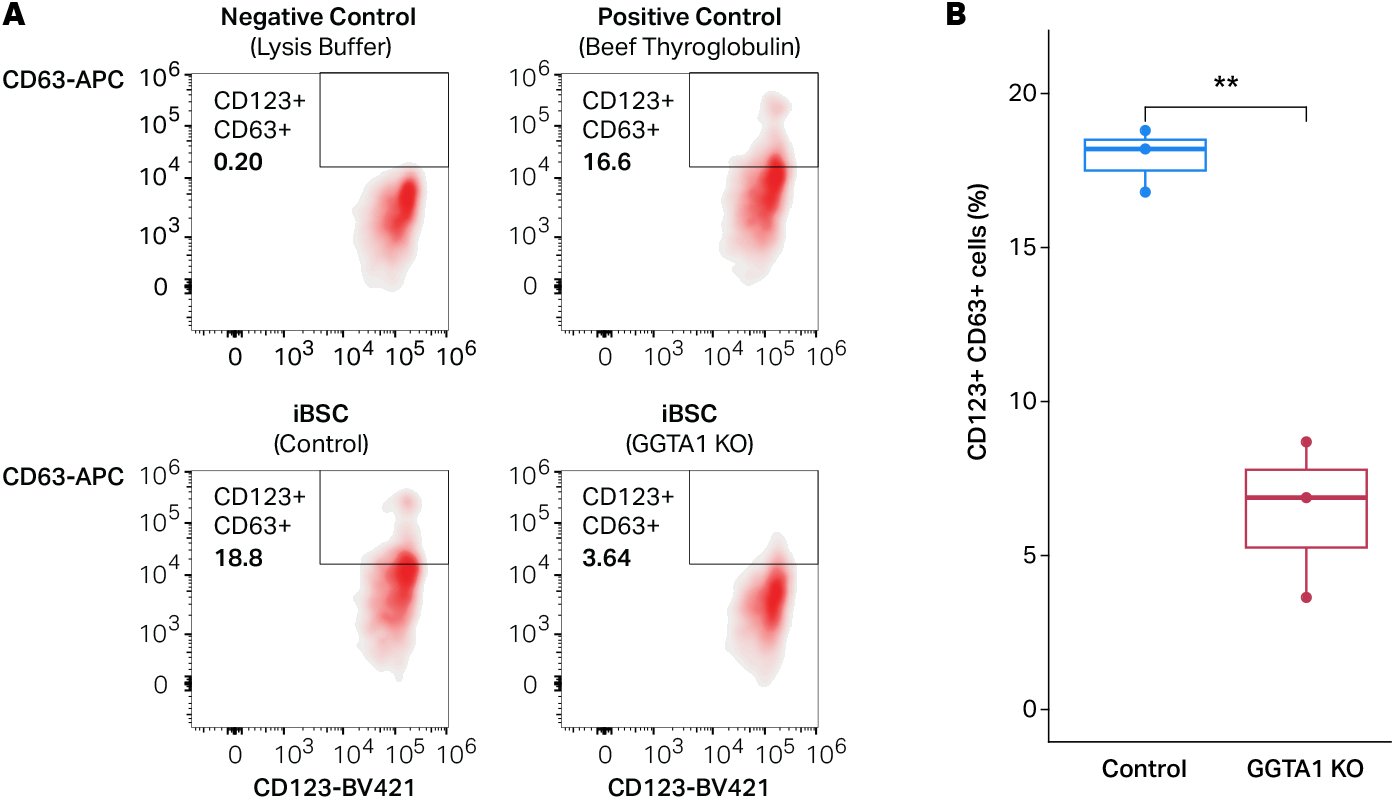
GGTA1 KO cell lysates induce reduced basophil activation in an indirect basophil activation test. **A**, Representative flow cytometry plots of human basophils following stimulation with lysis buffer negative control, beef thyroglobulin positive control, or lysates from control and GGTA1 KO iBSC clones. Basophil activation was measured as the percentage of CD123^+^ cells that were also CD63^+^. **B**, Quantification of basophil activation induced by lysates from control and GGTA1 KO clones. Each point represents an independent clone; bars show mean ± SD. Groups were compared using an unpaired two-tailed Welch’s t-test (p = 0.0083).

## Discussion

In this study, we demonstrate that targeted disruption of GGTA1 in immortalized bovine satellite cells can eliminate alpha-gal expression while preserving cellular morphology, differentiation capacity, and global transcriptional programs. Edited iBSCs exhibited significantly reduced basophil activation when tested using plasma from an alpha-gal–sensitized individual, indicating substantially reduced AGS-associated immune reactivity relative to unedited cells. Together, these findings establish a proof of concept that cultivated meat can be engineered to remove clinically relevant allergens while retaining key properties required for muscle cell expansion and differentiation.

Beyond addressing a specific unmet need in alpha-gal syndrome, this work highlights the broader potential of cultivated meat as a biotechnology platform. By enabling precise genetic modification of the cells that constitute meat, cultivated meat technologies create new opportunities to design food products that are not only more sustainable, but also potentially better tailored to human health needs. While additional work will be required to translate these findings into consumable products, our results represent an early step toward the development of alpha-gal–free cultivated beef and suggest how cell-based meat technologies may be used to address selected food-related health challenges.

Despite these promising results, several limitations should be considered. Although GGTA1 knock-out resulted in only a small number of differentially expressed genes relative to unedited controls, these subtle changes could impact meat production. While this observation warrants consideration, prior studies have successfully generated viable pigs (Phelps et al., 2003), cattle (Sendai et al., 2006), and other animals (McGregor et al., 2023; Tearle et al., 1996; Wei et al., 2022) from cells harboring comparable GGTA1 loss-of-function mutations, suggesting that disruption of alpha-gal synthesis is compatible with normal muscle development and organismal viability. Consistent with this, we did not observe detectable differences in morphology or differentiation-associated transcriptional programs between control and GGTA1 KO iBSCs, supporting the conclusion that myogenic capacity is largely preserved in the edited cells.

In addition, although GGTA1 knockout substantially reduced basophil activation, low-level residual activation was observed in some clones. One possible explanation is that alpha-gal is not the only non-human antigen present in bovine cells capable of engaging immune pathways in sensitized individuals. In addition, the use of cellular lysates, rather than processed and cooked meat, in the basophil activation assay may expose intracellular or non-food-relevant components that contribute to background activation. Notably, at least one knockout clone exhibited basophil reactivity approaching negative control levels, indicating that near-complete functional elimination of allergenicity is achievable in this system. Furthermore, pigs harboring analogous GGTA1 knockout mutations are currently used to produce “GalSafe” pork products (developed by Revivicor, Inc.), providing real-world evidence that loss of alpha-gal alone can be sufficient to mitigate allergenic risk in food products, despite low-level background immune reactivity observed in vitro. However, further testing will be needed to confirm safety.

Building on this proof of concept, several next steps will likely be important for advancing alpha-gal-free cultivated meat toward more translational and food-relevant applications. First, scale-up of GGTA1 knockout iBSCs under production-relevant culture conditions will be needed to assess whether alpha-gal elimination and preserved myogenic properties are maintained during large-scale expansion and differentiation. Second, while the present study focused on muscle progenitors, incorporation of additional cell types relevant to meat composition, such as adipogenic cells, will be important to determine whether alpha-gal elimination can be extended across multiple lineages and maintained in more complex, food-relevant cellular mixtures. Third, formulation of edited cells into structured, edible muscle tissue and processed food prototypes will help assess alpha-gal content and allergenicity in contexts that more closely resemble real-world consumption. Finally, further translation will require additional preclinical validation using food-relevant preparations and expanded patient-derived immune assays and, over the longer term, carefully designed and ethically appropriate human studies to evaluate safety.

Beyond alpha-gal syndrome, the strategy demonstrated here may be extensible to other food allergens or bioactive epitopes that pose health risks to specific populations. The ability to apply targeted genome editing to cultivated meat cell lines raises the possibility of multiplex engineering approaches that simultaneously address allergenicity, nutritional composition, and other health-relevant traits. As cultivated meat technologies mature, such applications could complement existing agricultural and biomedical strategies, positioning cell-based meat as a versatile platform for developing safer, healthier, and more nutritious food systems of the future.

## Materials and Methods

### Culture of immortalized bovine satellite cells (iBSCs)

Immortalized bovine satellite cells (passage 35) provided by David Kaplan’s laboratory (Tufts University) were expanded as previously described (Stout et al., 2023). Cells were cultured in growth medium consisting of DMEM+GlutaMAX (ThermoFisher, 10566016), 20% Fetal Bovine Serum (ThermoFisher, 26140079), 1X Antibiotic-Antimycotic solution (Thermo Fisher Scientific, 15240062), 2.5 ug/ml Puromycin (Thermo Fisher Scientific, A1113803) and 1 ng/mL human FGF-2 (PeproTech, 100-18B). When cells were seeded onto new culture surfaces 0.25 μg/cm2 iMatrix laminin-511 (Iwai North America, N-892021) was added to the media containing cells. Media was changed every 2-3 days and cells were harvested with 0.25% trypsin-EDTA (ThermoFisher, 25200056), and either passaged or cryopreserved in FBS with 10% dimethyl sulfoxide (DMSO) or 10% DMSO in growth media. For differentiating the cells, iBSCs were cultured until confluent and media was changed to DMEM+GlutaMAX containing 2% FBS for indicated days.

### CRISPR/Cas9 editing of Alpha-1-3-Galactosyltransferase (GGTA1) in iBSCs

A previously reported sgRNA targeting bovine GGTA1 was used in this study for gene knockout in iBSCs (Perota et al., 2019). The GGTA1 targeting guide RNA (Guide sequence-PAM (5’-3’) GAGAAAATAATGAATGTCAA-AGG is specific for start codon in exon 4. For transfection, iBSCs (passage 36) were trypsinized, washed with PBS and 0.8 million cells were resuspended in 100 µl of Primary Cell Nucleofector™ solution containing Supplement 1 (Lonza, V4XP-4012). The ribonucleoprotein complex was prepared by combining Alt-R™ S.p. Cas9 Nuclease V3 Cas9 (IDT, 1081058) with Alt-R CRISPR-Cas9 sgRNA (IDT) and PBS. Following incubation at room temperature for 15 minutes the RNP complex and Alt-R® Cas9 Electroporation Enhancer (IDT, 1075915) was added to the cells. The mixture was gently pipetted up and down and transferred to Nucleocuvette vessels (Lonza, V4XP-4012) and transfected using program ER-100 on a 4D-Nucleofector™ Core unit (Lonza). Cells were kept at room temperature for 8 minutes and then cultured in prewarmed antibiotic free growth media containing 20% FBS, 1 ng/mL human FGF-2 and iMatrix laminin-511. The next day media was replaced, cells were expanded and aliquots of the untransfected and transfected cells were used for DNA extraction and PCR. QuickExtract™ DNA Extraction Solution (Lucigen), was added to the cells and vortexed for a minute. Samples were then heated at 65 °C for 6 minutes, and 98 °C for 2 minutes and the extracted DNA solution was stored at -20 °C. PCR amplification was performed in a 20 µl reaction containing EconoTaq PLUS GREEN 2X Master Mix (Lucigen), 3 µl of DNA, 0.5 µM of each primer FW2: AGCATCTTTCACAACTCAGG and RV2: TGAGACATTAGGAACATGGC (Perota et al., 2019). PCR conditions included an initial denaturation at 94 °C for 2 minutes, 35 cycles of denaturation at 94 °C for 30 seconds, annealing at 60 °C for 30 seconds, and extension at 72 °C for 30 seconds, followed by a final extension at 72 °C for 7 minutes. Following amplification the PCR product was cleaned using the DNA clean and concentrator kit (Zymo Research) and submitted to GENEWIZ (South Plainfield, NJ) for Sanger sequencing with primers described above. The resulting sequencing data was analyzed using the Inference of CRISPR Edits (ICE) online tool from Synthego.

Following confirmation of GGTA1 editing in bulk transfected cells, single cell cloning of edited and control (untransfected) cells was attempted by diluting cells to 2 cells/ml and aliquoting 100 µl in each 96-well. A total of 95 wells for edited and 16 wells for control cells were seeded, however none of the cells developed into colonies. Edited and control (untransfected) cells were also seeded at a low cell density of 450 cells per 10 cm dish to get well separated clusters. Following culture, cell clusters were picked manually under a microscope (EVOS FL, ThermoFisher). Cell clusters were disrupted by pipetting and placed into 12-well plates for continued expansion and genetic analysis. Samples were screened using the DNA isolation and PCR protocol described above with some modification, PCR reaction volume was increased to 40 µl, and 1 µl of DMSO and 2 µl of colony DNA was added per reaction. Samples were sequenced and analyzed as described above. Based on sequencing results we proceeded with three control colonies and three GGTA1 KO colonies.

### RNA sequencing

To profile the transcriptome of control and GGTA1 KO cells before and after differentiation we conducted bulk RNA-sequencing. Three control and GGTA1 KO colonies were expanded in duplicate dishes to confluence in growth media containing FBS (20%).To assess differentiation, growth media was removed and replaced with DMEM containing 2% FBS for 3 days. Total RNA was extracted from cells before (d0) and after inducing differentiation (d3) with the RNeasy kit (Qiagen, 74104) according to the manufacturer’s recommendation. On-column DNase digestion was performed during the extraction, samples were quantified with the Qubit RNA High sensitivity assay kit (Thermo Fisher Scientific, Q32582) and RNA integrity number (RIN) was obtained using the Agilent TapeStation 4150. RIN values ranged from 9.3 to 10. RNA samples were shipped to GENEWIZ (South Plainfield, NJ) for library preparation (NEBNext Ultra II RNA Library Prep Kit for Illumina) with Poly (A) selection and 150-bp paired-end sequencing on a NovaSeq X Plus.

### Transcriptomic analysis

Quality control was performed with FastQC (v0.11.9) and MultiQC (v1.11)(Ewels et al., 2016). Low quality reads and adapters were trimmed with TrimGalore! (v0.6.7) (Krueger, 2019). Trimmed FASTQs were aligned to the BosTau9 genome with HISAT2 (v2.2.1) (Kim et al., 2019) and transcript-level quantifications were obtained with salmon (v1.10.0) (Patro et al., 2017). Transcript-level quantifications were summarized and converted to gene-level scaled transcripts in R (v4.5.0) with tximport and the txi file was imported into R using DESeqDataSetFromTXimport (Soneson et al., 2015). Differential gene expression analysis was performed with DESeq2 (v1.46.0) (Love et al., 2014). Before modeling, lowly expressed genes were omitted by requiring at least 10 counts in at least 6 samples. Genes were considered differential with an FDR-adjusted p value < 0.01 (Wald test) and an absolute log2 fold change > 1.

Gene Ontology (GO) enrichment analysis was done with gProfiler2 in R (Kolberg et al., 2020). Term enrichment for differentially up and down regulated gene sets were calculated separately for each comparison and the set of all expressed genes passing the filter for lowly expressed genes were used as background for the analysis. GO terms with a FDR-adjusted p value <0.05 were considered significantly enriched.

### Flow cytometric analysis and immunofluorescence staining for alpha-gal

For flow cytometric analysis, iBSCs were washed twice with PBS, placed in serum free media for an hour and washed again with PBS, then trypsinized and washed again with PBS. Cells from the control colonies and GGTA1 KO colonies were pooled, washed twice with PBS and stained with viability dye eFluor 450 according to manufacturer’s recommendation. Cells were then centrifuged, washed with PBS and fixed in 1% paraformaldehyde for 15 minutes. After washing with PBS, cells were resuspended in the 2% BSA blocking solution for an hour. Cells were incubated in mouse IgM monoclonal antibody solution against the alpha-gal epitope (Enzo Life Sciences ALX-801-090-1) at 1:5 dilution in PBS containing 1% BSA for 1 hour at room temperature. Cells were washed twice with PBS and incubated for an hour in anti-mouse IgM secondary antibody FITC (1:200; Thermo Scientific 31992). After a PBS wash cells were filtered with a 30 micron strainer to obtain single-cell suspension. In flow cytometry experiments, gates were determined based on using unstained cells as gating controls. Data was acquired on the Thermo Fisher Attune NxT (UNC Flow Cytometry Core Facility) and analyzed using FCS Express 8 software (De Novo Software, Glendale, CA, USA).

For immunofluorescent staining, approximately 60,000 control or knockout cells were seeded in a 24-well plate in DMEM, 20% FBS and 1X Antibiotic-Antimycotic solution, the next day, cells were washed three times with PBS and incubated for an hour in serum free media (DMEM+GlutaMAX) followed by three washes with PBS. Cells were fixed in 4% formaldehyde (Thermo Scientific 28908) for 20 min. After a PBS wash, cells were blocked for an hour in 2% BSA and then incubated in the alpha-gal antibody solution (1:5; Enzo Life Sciences, ALX-801-090-1) in 1% BSA for 1 hour at room temperature. Cells were washed twice with PBS and incubated for an hour in anti-mouse IgM secondary antibody FITC (1:200; Thermo Scientific, 31992). Following two washes in PBS, cells were mounted with Slowfade Gold Antifade Mountant containing DAPI (Thermo Scientific, S36939).

### Protein extraction

Control and knockout clones were grown in 10 cm dishes, washed with PBS and media changed to serum free media (DMEM+GlutaMAX) for 2h. Following two PBS washes, lysis buffer (Cell signaling technology, 9803) containing PMSF (1mM) was added to the dishes and incubated on ice for 5 minutes. Plates were scraped, lysate collected and rotated at 4C for 30 minutes and then frozen at -70C. Thawed lysates were sonicated and centrifuged 12,000g for 10 min at 4C. Supernatant was collected and stored at -70C. The protein concentration was determined using the Micro BCA Protein Assay kit (Fisher Scientific, PI23235). Protein samples were diluted to 1µg/µl and 5 µg of lysate was used per basophil activation test reaction.

### Indirect basophil activation test

Basophils from healthy individuals’ PBMCs were isolated using a Ficoll-Paque density gradient (GE Healthcare, Chicago, IL). IgE was stripped from the basophils using a cold lactic acid buffer (13.4 mM lactic acid, 140 mM NaCl, 5 mM KCl) for 10 minutes on ice. Cells were washed with RPMI-1640 and then sensitized with the 50% plasma of an alpha-gal allergic subject in RPMI1640 cell culture media (Corning Cell-Gro, Manassas, VA, USA) containing 1 ng/mL IL-3 (R&D Systems, Minneapolis, MN, USA) overnight at 37°C with 5% CO2 (Sharma et al., 2024). Basophils were stimulated for 30 minutes with basophil activation media alone (RPMI plus 2 ng/mL IL-3) or activation media plus test reagents (cellular lysate from control and knockout clones), and stimulation was stopped by adding EDTA (2.5 mM). Basophil media (BM) and lysis buffer used for protein extraction were used as negative controls. Rabbit anti-human IgE antibody (1 µg / ml; Bethyl Laboratories Inc., A80-109A), cetuximab (1 µg/ml; Eli Lilly and Company, Indianapolis, IN), and beef thyroglobulin (10 µg/ml; Sigma-Aldrich, St. Louis, MO) were used as positive controls. Cells were stained for flow cytometry utilizing fluorescently labeled antibodies, including the basophil marker CD123-BV421 (BD Biosciences, San Jose, CA), the basophil activation marker CD63-APC (BioLegend, San Diego, CA), HLA-DR-PerCp-Cy5.5 (ThermoFisher Scientific, Waltham, MA), and FITC lineage cocktail 1 (comprising CD3, CD14, CD16, CD19, CD20, CD56; BD Biosciences, San Jose, CA). The samples were run on an Attune NxT flow cytometer (ThermoFisher Scientific), and data were analyzed using FlowJo v10.9.0. Percentages of CD63-positive basophils (lineage-1-, HLA-DR-, CD123+, CD203c+) were determined and used as a marker of basophil activation. Statistical analysis was performed using GraphPad Prism version 10.6.1. Welch’s t-test was used to compare two groups.

## Data Availability

All raw RNA sequencing data supporting the findings of this study are available in the NCBI Gene Expression Omnibus (GEO; https://www.ncbi.nlm.nih.gov/geo/) under the accession GSE330550.

## Code Availability

All analysis code used in this study is available in a git repository at https://github.com/grkenney/AGAL.

## Artificial Intelligence Statement

During the preparation of this work the author(s) used ChatGPT in order to improve clarity. After using this tool/service, the author(s) reviewed and edited the content as needed and take(s) full responsibility for the content of the published article.

## Author Contribution

**Susan D’Costa:** Formal analysis, Investigation, Methodology, Visualization, Writing – original draft.

**Shailesh K. Choudhary:** Formal analysis, Investigation, Methodology, Visualization, Writing – original draft.

**Grace E. Kenney** Formal analysis,: Data curation, Investigation, Methodology, Visualization, Writing – original draft.

**Jacqueline Shine:** Formal analysis, Visualization.

**Brian Diekman:** Supervision.

**Scott P. Commins:** Conceptualization, Funding acquisition, Project administration, Resources, Supervision, Writing – review & editing.

**Douglas H. Phanstiel:** Conceptualization, Funding acquisition, Project administration, Supervision, Writing – review & editing.

## Acknowledgements

We thank Andrew Stout (Kaplan lab, Tufts University) for providing iBSCs, Erika Deoudes for data visualization, illustration, and typesetting. We also acknowledge the UNC Flow Cytometry Core Facility for technical assistance with alpha-gal flow cytometry, the UNC Flow Cytometry Core Facility (RRID:SCR_019170) is supported in part by P30 CA016086 Cancer Center Core Support Grant to the UNC Lineberger Comprehensive Cancer Center. Research reported in this publication was supported in part by the North Carolina Biotech Center Institutional Support Grant 2017-IDG-1025 and by the National Institutes of Health 1UM2AI30836-01. The content is solely the responsibility of the authors and does not necessarily represent the official views of the National Institutes of Health. Graphical abstract created with BioRender.com.

